# Memory B cell Development in Response to mRNA SARS-CoV-2 and Nanoparticle Immunization in Mice

**DOI:** 10.1101/2025.10.01.679787

**Authors:** Marie Wiatr, Zijun Wang, Marie Canis, Brianna Hernandez, Anna Gazumyan, Gabriela S. Silva Santos, Sadman Shawraz, Songhee Lee, Paul D. Bieniasz, Theodora Hatziioannou, Frauke Muecksch, Michel C. Nussenzweig

**Affiliations:** Laboratory of Molecular Immunology, The Rockefeller University; New York, NY 10065, USA; Laboratory of Retrovirology, The Rockefeller University; New York, NY 10065, USA; Howard Hughes Medical Institute, The Rockefeller University, New York, NY, USA; Department of Infectious Diseases, Virology, Medical Faculty Heidelberg, Heidelberg University; 69120 Heidelberg, Germany

**Keywords:** Nanoparticle vaccine, mRNA vaccine, SARS-CoV-2, memory B cell, neutralizing antibodies

## Abstract

Nanoparticle immunogens excel at rapidly inducing high levels of circulating antibodies and are being deployed as part of several novel vaccines. However, their ability to elicit memory B cell responses is less well understood. Here we compared serologic and memory B cell responses to prime boost vaccination with either SARS-CoV-2 Wuhan-Hu-1 mRNA vaccine, or protein nanoparticles: SARS-CoV-2 B.1.351 homotypic containing a single receptor binding domain (RBD); (homotypic beta) or a combination of different Sarbecovirus RBDs (mosaic 8b), respectively. The memory B cells elicited by the 3 vaccine regimens showed closely related antibody sequences, similar levels of somatic mutation and clonal diversity. The breadth of serologic responses elicited by the mosaic nanoparticles were comparable to the homotypic nanoparticle and superior to the mRNA vaccine for some mismatched strains. However, serum neutralizing titers to SARS-CoV-2 were highest after mRNA vaccination. The three vaccines elicited memory B cells that produced antibodies specific to a broad range of epitopes on the RBD that differed in a way that may reflect epitope masking. Monoclonal antibodies derived from memory B cells elicited by the mosaic 8b nanoparticle showed greater breadth against a panel of SARS-CoV-2 variants and SARS-CoV.

**Significance Statement:** Nanoparticle vaccines are promising next-generation vaccine candidates, yet their capacity to generate durable memory B cell responses remains incompletely understood. We compared immune responses following SARS-CoV-2 mRNA, homotypic beta nanoparticle, or mosaic 8b nanoparticle vaccination in mice. Serum antibody neutralizing responses against a panel of SARS-CoV-2 variants elicited by an mRNA vaccine were equivalent or superior to those elicited by mosaic 8b nanoparticle vaccines. However, the monoclonal antibodies derived from memory B cells elicited by the mosaic 8b nanoparticle showed better neutralizing breadth against heterologous pseudoviruses. These findings highlight individual strengths of mRNA and nanoparticle vaccines and show that mosaic 8b nanoparticle immunogens can enhance the breadth of memory B cell-derived antibodies.

## Introduction

Vaccine efficacy depends on both cellular and humoral immunity. Whereas circulating antibodies can prevent infection, memory T cells and B cells respond rapidly to breakthrough infection and can limit disease progression. Long-lived circulating antibodies are produced by plasma cells that are selected by a mechanism which favors high affinity antibody production (1-6). In contrast, memory B cell selection is governed by a distinct mechanism that produces cells with diverse affinity (7-9). As a result, the memory compartment in immunized mice includes B cells producing antibodies that are cross-reactive to antigens that are related but not identical to the immunogen (8-10).

The recent coronavirus pandemic enabled comparative studies of human memory B cell development in response to SARS-CoV-2 infection and/or vaccination (11-15). Several different vaccines were studied and compared to natural infection for their ability to elicit memory B cells (16-20). Overall, the data indicated that mRNA vaccination was most effective in producing memory B cells and preventing serious disease in individuals with breakthrough infections (21, 22).

Humans receiving mRNA vaccines developed a memory B cell compartment that evolved increased breadth and potency over time (15, 23-25). Thus, a significant fraction of memory B cell antibodies elicited by an mRNA vaccine encoding the original Wuhan-Hu-1 Spike also neutralized emerging variants (23, 24, 26). Increasing breadth after SARS-CoV-2 mRNA vaccination was attributed to changes in epitope targeting directed by antibody masking (27). The cross-reactive memory B cells likely contributed to the observation that breakthrough infections with variants of concern were less likely to result in serious disease in vaccinated individuals (28-30).

Coronaviruses are found in many ecological niches and are particularly common in bat species, which make up 20% of all mammals (31). Therefore, future coronavirus spillovers into the human population are likely (32). Memory B cell responses to the current SARS-CoV-2 mRNA vaccines are not universally effective against distantly related coronaviruses making it imperative to explore vaccine strategies that protect against a broader group of these viruses. One of the vaccine concepts that appears to broaden the immune response involves the use of mosaic nanoparticles that display a mixture of different coronavirus receptor binding domains (RBDs), the primary target of neutralizing antibodies (33-35).

Nanoparticle vaccine concepts are also being tested for Influenza virus and HIV-1 (36-40). In all cases, nanoparticle vaccines elicit protective titers of circulating antibodies. However, the impact of nanoparticle vaccines on B cell memory development has not been evaluated in comparison to mRNA vaccination (33, 34, 41-44). Here we examine the development of serum antibody and B cell memory responses to homotypic beta and mosaic 8b RBD nanoparticle vaccination and compare them to responses elicited by SARS-CoV-2 mRNA vaccines in mice.

## Results

To examine memory B cell responses to candidate nanoparticle vaccines we primed C57BL/6J and BALB/c mice with SpyCatcher003-mi3 nanoparticles coupled to SpyTag003-RBDs (33) (Fig. 1A, B). The mice were boosted 28 days after the prime with the same formulation (Fig. 1A, B).

**Figure 1.**
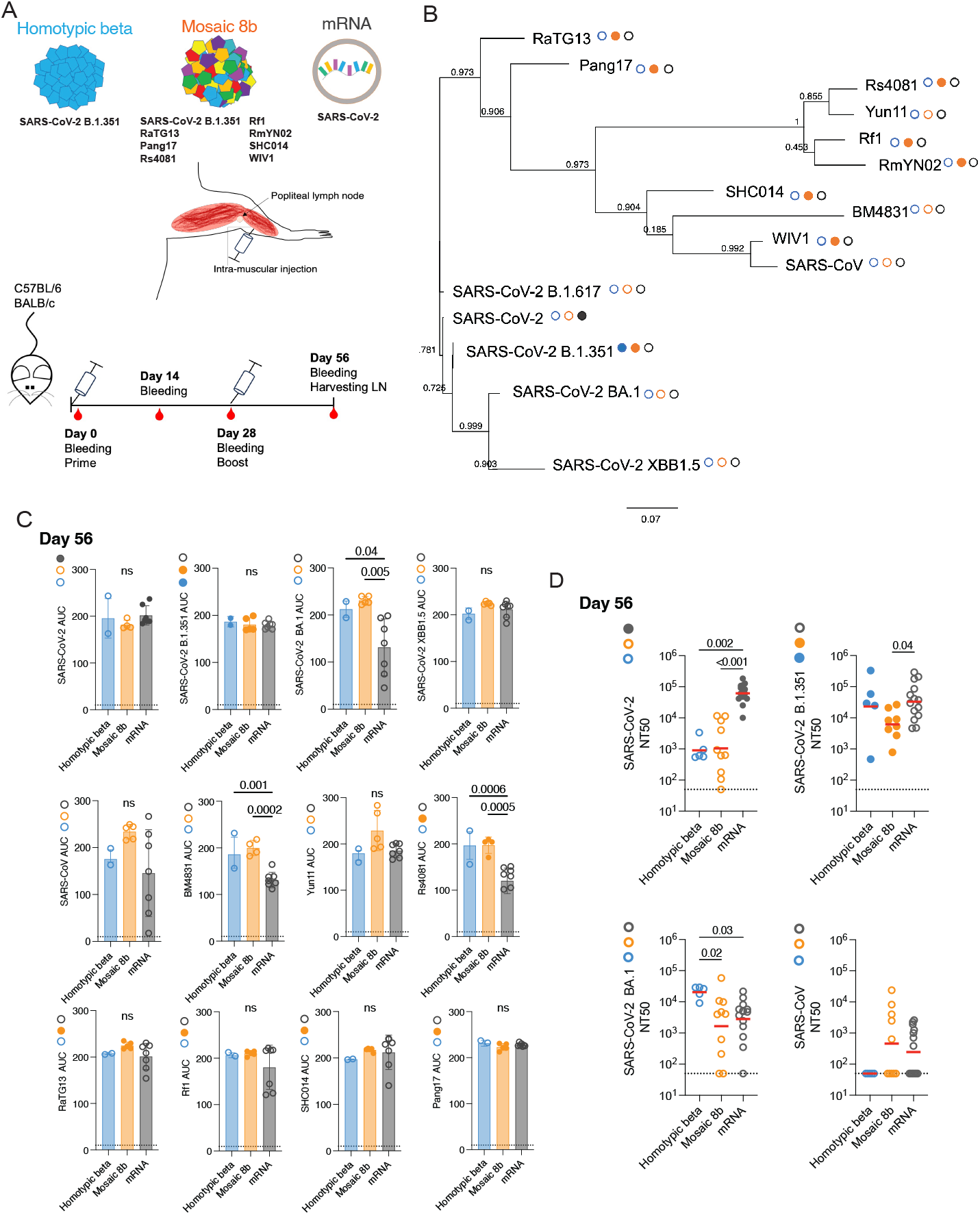
Plasma binding and neutralizing activity. (A) Schematic representation of the vaccines, and immunization protocol. (B) Phylogenetic tree showing amino acid proximity of SARS-CoV-2 RBD with other Sarbecoviruses. Filled circles next to strain names designate strains that are represented in the vaccine (matched) and unfilled circles designate strains that are not represented in the vaccine (mismatched). (C) Graphs show plasma IgG binding activity for samples obtained from C57BL/6 mice as normalized area under the curve (AUC) values measured by ELISA for: SARS-CoV-2, SARS-CoV-2 B.1.351, SARS-CoV-2 BA.1, SARS-CoV-2 XBB1.5, SARS-CoV, BM4831, Yun11, Rs4081, Pang17, RaTG13, Rf1, and SHC014 RBDs on day 56. Solid colored dots indicate the presence of the RBD on the immunogen [homotypic beta (blue) mosaic 8b (orange) mRNA (gray)]. Filed circles represent presence of the RBD in the immunogen. The dotted black line represents negative control. Each dot represents one animal. P values were calculated using an ANOVA test. (D) Plasma neutralizing titers for SARS-CoV-2, SARS-CoV-2 B.1.351, SARS-CoV-2 BA.1 and SARS-CoV for samples obtained from C57BL/6 and Balb/c mice (homotypic beta: n=3 and n=2, mosaic 8b: n=5 and n=5, mRNA: n=7 and n=7 for C57BL/6 and Balb/c mice, respectively). Plasma NT50s for homotypic (blue), mosaic 8b (orange) and mRNA (gray) measured on day 56. Each dot represents one animal. The horizontal bar represents the geometric mean. All experiments were performed at least twice. P values were calculated using Kruskal-Wallis test followed by Dunn’s multiple comparison test. Only P values > 0.05 are shown on the graph.

The nanoparticles carried either SARS-CoV-2 B.1.351 RBD (homotypic beta), or a combination of eight different RBDs corresponding to SARS-CoV-2 B.1.351, RaTG13, Pang17, Rf1, Rs4081, SHC014, RmYN02, and WIV1 (mosaic 8b nanoparticles) (Fig. 1A, B and Table S1) (33). This collection of RBDs includes sequences representative of sarbecoviruses corresponding to clades 1, 2, 1/2 and 3, covering a broad spectrum of genetic diversity within this subgenus (33). A separate cohort of control mice was primed and boosted 28 days after the prime with the original monovalent SARS-CoV-2 mRNA vaccine encoding a pre-fusion stabilized Wuhan-Hu-1 spike protein (45) (Fig. 1A, C).

### Plasma binding activity

Antibody titers to RBD were measured 14 days after the prime, shortly before the boost on day 28, and 4 weeks after the boost on day 56 (Fig. 1C and Fig. S1A, B). Twelve different RBDs were used to assess binding by ELISA: SARS-CoV-2 which is encoded by the mRNA vaccine; SARS-CoV-2 B.1.351 (46) present on homotypic beta and mosaic 8b particles; Rs4081, Pang17, RaTG13, Rf1, SHC014 found exclusively on the mosaic 8b nanoparticle; and 5 RBDs not found on any of the immunogens: SARS-CoV-2 XBB1.5, SARS-CoV-2 BA.1, BM4831, SARS-CoV and Yun11 (Fig. 1B, D and Fig. S1A, B (47-49)).

Two weeks after the prime, there were no significant differences in serum binding titers to 5 of the 12 RBDs tested between the vaccination groups (Fig. S1A). The exceptions were SARS-CoV against which the mosaic 8b nanoparticle and mRNA produced higher binding antibodies than homotypic beta and BM4831, Yun11, Rs4081, Pang17, RaTG13, and Rf1, where mosaic 8b was best (Fig. S1A). These differences were slightly less apparent 28 days after the prime (Fig. S1B).

In all 3 vaccination groups, boosting produced higher antibody titers (Fig. 1C). After the boost there were no significant differences between homotypic beta and mosaic 8b nanoparticle immunization (Fig. 1C); however, both were significantly better than the mRNA vaccine against SARS-CoV-2 BA.1, BM4831 and Rs4081 (Fig. 1C). Thus, the breadth and overall amount of serologic RBD binding activity was better after nanoparticle than mRNA vaccination.

### Plasma neutralization activity

To measure serum neutralizing activity, we used HIV-1-based virions carrying a Nanoluciferase reporter pseudotyped with either SARS-CoV-2 WT, SARS-CoV-2 B.1.351, SARS-CoV-2 BA.1 or SARS-CoV spikes (50) (Fig. 1D, Fig. S1C, D). mRNA-vaccinated mice showed significantly higher neutralizing activity against SARS-CoV-2 than either of the nanoparticle vaccines after both prime and boost (Fig. 1D and Fig. S1C, D). After the boost, mRNA-elicited titers against SARS-CoV-2 B.1.351 were significantly higher than after mosaic 8b vaccination (Fig. 1D) and homotypic beta nanoparticle-vaccinated mice showed significantly higher neutralizing titers against SARS-CoV-2 BA.1 than mice vaccinated either with mRNA or mosaic 8b nanoparticles. Furthermore, the amount of SARS-CoV neutralizing activity elicited by the mosaic 8b after boosting was not significantly different from the SARS-CoV-2 mRNA vaccination. (Fig. 1D, Fig. S1C, D).

### Memory B cells

Memory B cells are responsible for anamnestic serologic responses (53). These cells are selected by different mechanisms than serum antibody-producing plasma cells (51, 52). To examine the antigen-binding memory B cell compartment in the immunized mice we purified lymph node B cells that bind to SARS-CoV-2 RBD 4 weeks after the boost (Day 56) and cloned their antibody genes (Fig. 2 and Fig. S2A, B). We selected SARS-CoV-2 for memory B cell capture because this RBD and SARS-CoV-2 B.1.351 had the same serological titers at day 56 and no significant difference in the 3 groups of vaccines. The absolute number of B cells binding to RBD was similar in the three groups of immunized mice (Fig. 2A and Fig. S2B). 309 Ig heavy and light chain pairs were obtained, many of which were found in expanded clones (Fig. 2B and Table 2). These antibodies from homotypic beta nanoparticle, mosaic 8b nanoparticle, and mRNA vaccination showed closely related V+J sequences (Fig. 2C). In addition, the antibodies had similar levels of diversity and CDR3 length (Fig. S2C, D). Finally, the relative amount of clonal expansion was similar for all 3 immunization regimens, as was the amount of somatic mutation (Fig. 2B, D).

**Figure 2.**
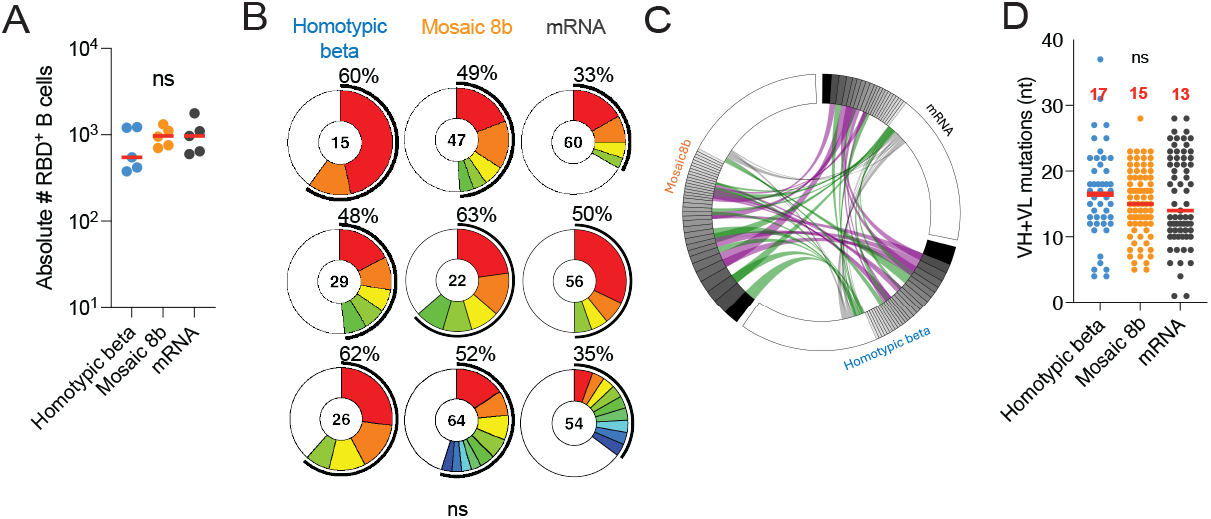
Memory B cell antibodies. (A) Absolute number of RBD binding B cells in homotypic beta- (n=5 in blue), mosaic 8b- (n=5 in orange) and mRNA- (n=5 in gray) vaccinated mice obtained from draining lymph node on day 56. (B) Pie charts showing the proportional clonal representation of antibodies obtained from the memory B cells in (A). The number indicates the number of sequences. The black outline and percentage show the frequency of clonally expanded B cells. Colors represent individual clones. (C) Circus plot shows V+J antibody gene segments shared among mice immunized with the different immunogens. Purple shows antibodies shared within clones. Green shows antibodies shared from clones to singles and grey shows shared single antibodies between the 3 immunizations. (D) Nucleotide somatic hypermutations (SHM) in the heavy and light chains for all sequences shown in (B) for homotypic beta (blue), mosaic 8b (orange) and mRNA (gray) immunization groups. The red line represents the median value. Statistical significance was determined using an ANOVA test.

### Monoclonal antibody binding activity

To further characterize the memory response, we expressed 211 randomly selected monoclonal antibodies: 55 from homotypic beta-, 81 from mosaic 8b-, and 75 from mRNA-immunized mice (Table 2). Binding activity was measured by ELISA against SARS-CoV-2 RBD (only present on the mRNA), SARS-CoV-2 B.1.351 RBD (present on homotypic beta and mosaic 8b nanoparticle), Rs4081 and Pang17 RBDs (only present on mosaic 8b), and SARS-CoV-2 BA.1, SARS-CoV-2 XBB1.5, SARS-CoV-2 B.1.617, Yun11, BM4831 and SARS-CoV RBDs (absent from all vaccines; Fig. 1B and 3A). For all RBDs tested, binding by the mosaic8b-elicited antibodies was similar to those elicited by the mRNA vaccine, except for SARS-CoV RBD for which binding by mosaic 8b-elicited antibodies were better (Fig. 3A). The mosaic 8b nanoparticle antibodies were significantly better binders than homotypic beta-elicited antibodies to SARS-CoV- 2 B.1.617, Yun11, Rs4081, BM4831 and SARS-CoV (Fig. 3A).

**Figure 3.**
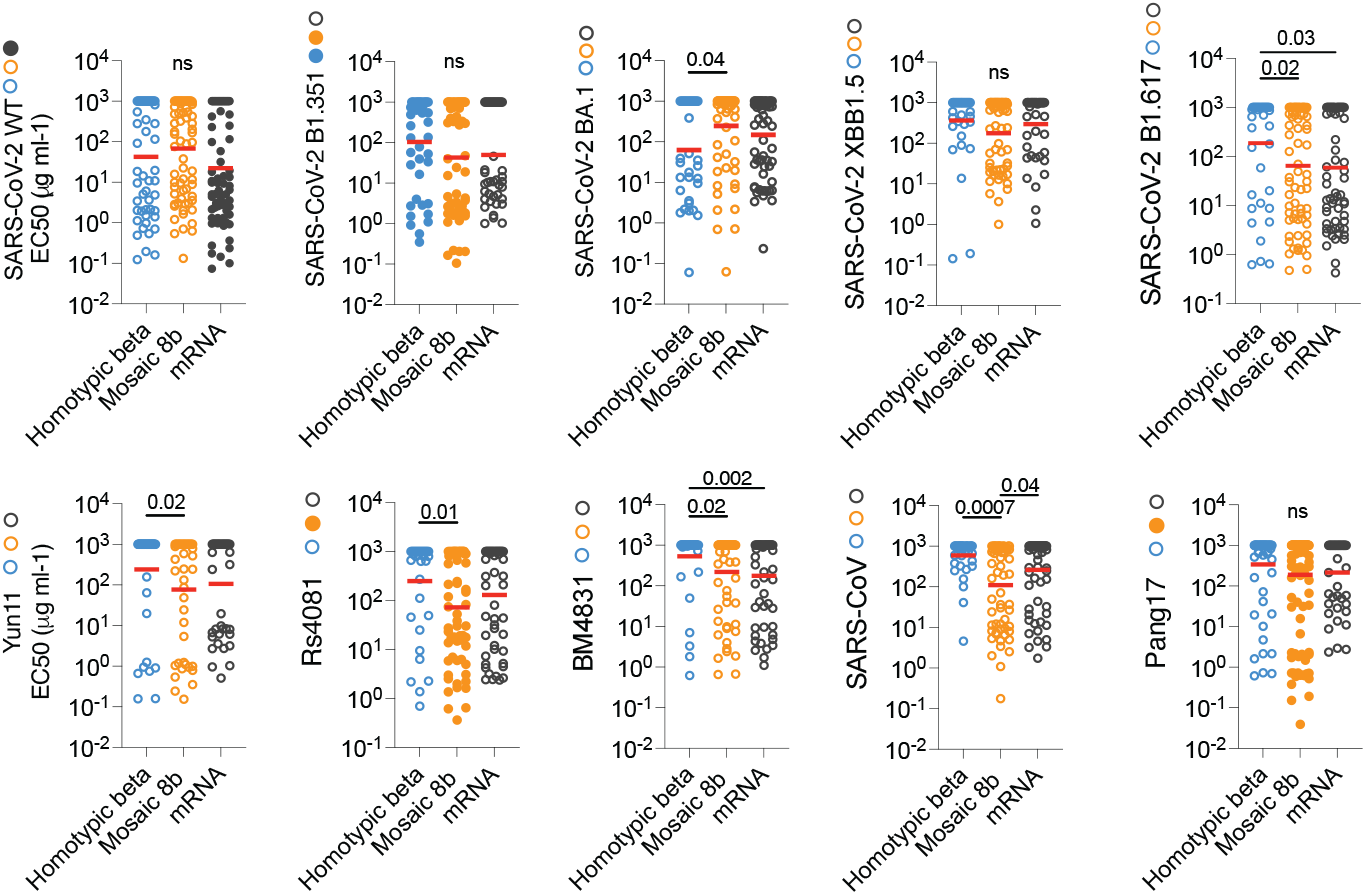
Binding activity of monoclonal antibodies. (A) Graphs show monoclonal antibody binding activity measured by ELISA (EC50s) for: SARS-CoV-2, SARS-CoV-2 B.1.351, SARS- CoV-2 BA.1, SARS-CoV-2 XBB1.5, SARS-CoV-2 B.1.617, Yun11, Rs4081, BM4831, SARS-CoV, Pang17 RBDs. Filled colored dots show presence of the respective RBD on the immunogen [homotypic beta (blue) mosaic 8b (orange) mRNA (gray)]. Each dot represents one monoclonal antibody. Red lines indicate geometric mean. Statistical significance was determined using an ANOVA test. Only P values> 0.05 are shown on the graph.

### Monoclonal antibody neutralization activity

To determine the neutralizing activity of the memory antibodies we tested all 144 antibodies with detectable binding activity to SARS-CoV-2 or SARS-CoV-2 B.1.351 against viruses pseudotyped with SARS-CoV-2, SARS-CoV-2 B.1.351, SARS-CoV-2 BA.1, SARS-CoV-2 XBB1.5, and SARS-CoV spikes (31 from homotypic beta nanoparticle-, 47 from mosaic 8b nanoparticle-, and 66 from mRNA-immunized mice). Overall, 61%, 74%, and 62% of the binding antibodies cloned from memory B cells obtained from homotypic beta, mosaic 8b nanoparticle, and mRNA vaccinated mice, respectively, showed measurable neutralizing activity to SARS-CoV-2 or SARS-CoV-2 B.1.351 with an IC50 of less than 1 µg/ml (Fig. 4A, and Fig. S3A).

**Figure 4.**
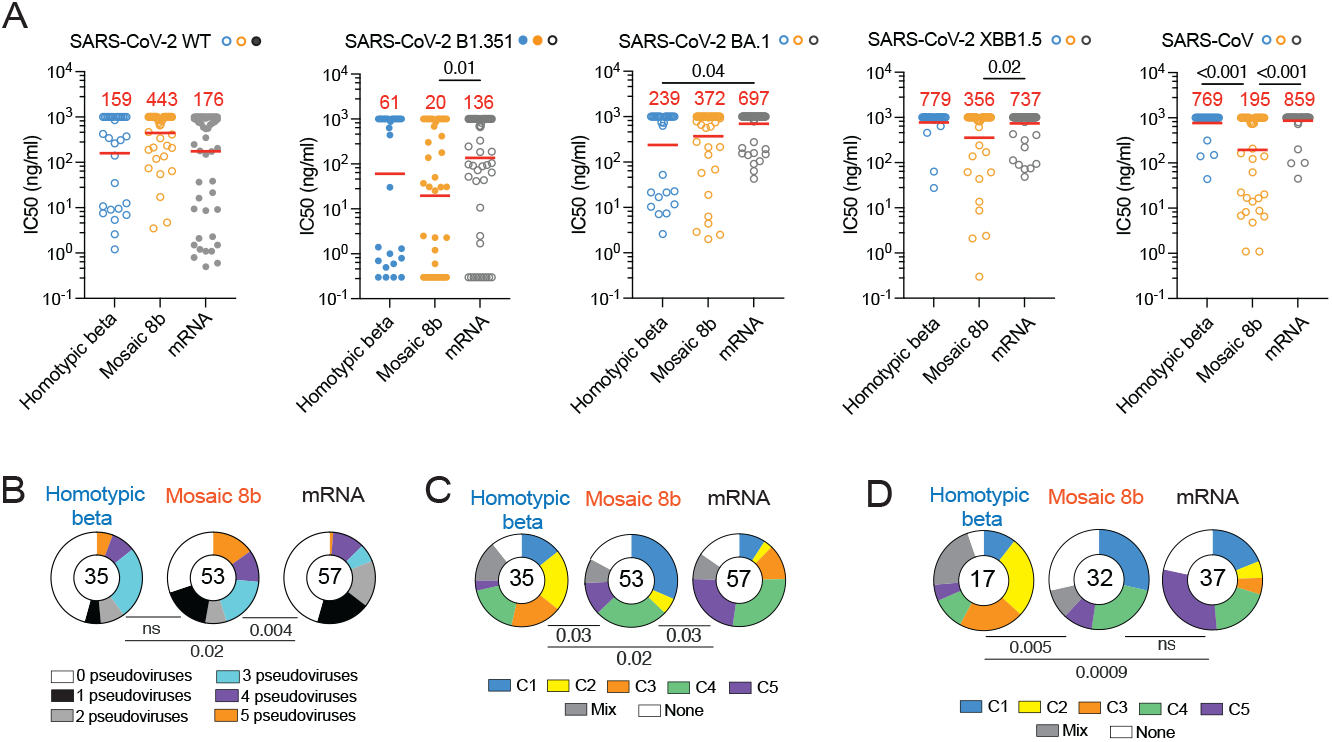
Neutralizing activity and target epitopes of monoclonal antibodies. (A) SARS-CoV-2, SARS-CoV-2 B.1.351, SARS-CoV-2 BA.1, SARS-CoV-2 XBB1.5 and SARS-CoV pseudovirus neutralization activity. IC50 values in ng/ml for the indicated pseudoviruses. The red line shows the geometric mean indicated in red. Statistical significance was determined using the Kruskal-Wallis test with subsequent Dunn’s multiple-comparisons test. (B) Pie charts depict the fraction of antibodies that neutralize 0 (white), 1 (black), 2 (gray), 3 (blue), 4 (purple), 5 (orange) pseudoviruses with an IC50<1000 ng/ml. The number indicates the number of antibodies tested. Statistical significance was determined using the two-sided Fisher’s exact tests. (C) Pie chart represents epitope target class for all monoclonal antibodies with binding activity to SARS-CoV-2 RBD determined by competition BLI. The number indicates the number of antibodies tested. (D) Pie chart represents epitope target class for all monoclonal antibodies that neutralize more than 1 pseudovirus. The number indicates the number of antibodies tested. Statistical significance in B- D was determined using two-sided Fisher’s exact tests.

The fraction of SARS-CoV-2 neutralizers obtained from mRNA immunized mice is consistent with the fraction obtained from vaccinated humans (19). The geometric mean of the neutralizing activity was 159, 443, and 176 ng/ml for the homotypic beta, mosaic 8b nanoparticles, and mRNA memory antibodies, respectively, with no significant difference between the mosaic8b- elicited antibodies and either homotypic beta- or mRNA vaccine-elicited antibodies (Fig. 4A and Fig. S3A). Although the number of antibodies is limited, these trends remain the same when considering neutralizing antibodies alone (Fig. S3B). Mosaic 8b-derived antibodies had significantly better neutralizing activity against SARS-CoV-2 B.1.351 and SARS-CoV-2 XBB1.5 than the mRNA vaccine-derived antibodies (Fig. 4A and Fig. S3A). As might be expected, antibodies derived from mice vaccinated with the mosaic 8b nanoparticle were significantly better neutralizers of SARS-CoV than the other 2 immunogens (Fig. 4A, Fig. S3A). However, there were no statistically significant differences between the memory antibodies elicited by the mosaic 8b nanoparticles vs. the other two for neutralization of SARS-CoV-2 BA.1-pseudotyped viruses (Fig. 4A, Fig. S3A).

The breadth of neutralizing activity was evaluated by comparing the ability of any one antibody to neutralize 0 to 5 of the pseudotyped viruses (Fig. 4B and Fig. S3A). The distribution of antibodies neutralizing 0 to 5 strains elicited by homotypic beta and mosaic 8b nanoparticles was not significantly different (Fig. 4B and Fig. S3A). Moreover, both sets of nanoparticle vaccine antibodies showed increased overall breadth of neutralization when compared to the mRNA vaccine (Fig. 4B and Fig. S3A).

### Epitopes Targeted by Monoclonal Antibodies

Memory antibodies elicited after the initial mRNA prime and boost in humans typically target Class 1 and 2 epitopes that overlap with the ACE2 binding site of the RBD (53, 54). After the 3rd dose, antibody-mediated epitope masking shifts the immune response to target more conserved features of the RBD between Class 3, 4 and 5 domains (23). To determine the specificity of the memory antibodies elicited by nanoparticle and mRNA prime-boost vaccination in mice, we performed biolayer interferometry (BLI) experiments in which a preformed antibody-RBD immune complex was exposed to a second monoclonal (reference) with predetermined specificity (55).

The 5 reference antibodies, C022, C144, C135, C837, and C5078, bind to Class 1, 2, 3, 4, 5, respectively (Fig. 4C and Fig. S4 (56)). Antibodies that bound to Class 1 or 2 (blue and yellow) were more prominent in the memory antibodies obtained from homotypic beta and mosaic 8b nanoparticle- than mRNA- immunized mice (Fig. 4C and Fig. S4). In contrast, the memory antibodies elicited by mRNA immunization were significantly different from the nanoparticle antibodies in that they primarily targeted the Class 3, 4 and 5 domains of the RBD (Fig. 4C and Fig. S4).

Broader antibodies with neutralizing activity against more than 1 pseudovirus also differed significantly between the groups. The combined class 3/4/5 antibodies that target the more conserved region of the RBD were more highly represented among neutralizers elicited by the mRNA vaccine than among those obtained from homotypic beta and mosaic 8b vaccination (Fig. 4D).

In conclusion, the mRNA vaccine elicits memory B cells producing antibodies that are less broad than those elicited by homotypic beta or mosaic 8b nanoparticle vaccines.

## Discussion

We compared serologic and memory B cell responses elicited by two different nanoparticle vaccines to those elicited by an mRNA vaccine in mice. Consistent with prior analysis (33), the binding activity of antibodies in serum elicited by the mosaic 8b nanoparticle vaccine carrying a diverse collection of coronavirus RBDs was somewhat broader than those elicited by an mRNA vaccine. When comparing serum neutralizing activity, the mRNA-vaccine elicited significantly higher neutralizing titers to its cognate antigen, SARS-CoV-2, and titers equivalent to the nanoparticles against their cognate antigen, SARS-CoV-2 B.1.351.

Nanoparticles can resemble pathogens in that they display multiple copies of antigen on their surface (57). These repeating units can cross-link antigen-specific receptors on B lymphocytes and produce rapid antibody responses that help clear the pathogen (58, 59). Consistent with this idea, mosaic 8b nanoparticle vaccination elicited a rapid initial antibody response that was protective against infection with either SARS-CoV-2 or SARS-CoV in experimental animals (33, 34, 41, 43, 44).

Memory B cell responses produced by nanoparticle vaccination showed similar levels of somatic mutation to those elicited by mRNA vaccination, suggesting that memory cells that develop after nanoparticle and mRNA vaccination do so after equivalent numbers of cycles of mutation and selection in germinal centers (60). However, the relative fraction of B cells producing anti-RBD neutralizing antibodies was slightly higher for animals receiving the mRNA vaccine. This may in part be due to the mRNA vaccine resulting in expression of a complete stabilized spike in its native conformation, whereas the RBD displayed on the nanoparticle is devoid of other components of the spike that may block non-neutralizing epitopes on the RBD. In addition, the relative effects of dosing cannot be excluded, and how these results obtained in mice might translate to humans is difficult to predict.

Germinal center responses are not required for memory B cell formation, but memory cells emerging from germinal centers are more likely to carry somatic mutations (60, 61). The mutations diversify the germinal center-derived memory B cells resulting in a collection of cells some of which produce highly specific antibodies and others that may also be able to recognize related antigens (8-10). Thus, germinal center diversification and selection produce memory B cells that can be called upon to respond to and neutralize related antigenic variants (11, 12, 23, 24). In addition to T cell responses to conserved epitopes, the diversified memory B cell compartment elicited by mRNA vaccination likely contributed to protection from the serious consequences of infection in individuals who experienced breakthrough infections with SARS-CoV-2 variants after mRNA vaccination (28-30).

Although all 3 vaccines appeared to elicit comparable germinal center responses, the epitopes targeted by memory B cells differed between nanoparticle and mRNA vaccines. In both humans and mice, the initial response to mRNA vaccination is dominated by antibodies targeting the more exposed Class 1 and 2 epitopes with a shift to less accessible Class 3,4,5 epitopes after a boost due to epitope masking (27). Compared to this, the fraction of Class 3,4,5 antibodies is reduced after a nanoparticle boost. In addition to the topological differences in the antigen presented by the immunogens, the higher overall level of Class 3,4,5 antibodies that develop after mRNA immunization may be due to initial epitope masking by Class 1 and 2 antibodies (27).

In summary, both SARS-CoV-2 mRNA and nanoparticle vaccines elicit germinal center reactions that promote production of a diverse memory B cell compartment enriched in neutralizing antibodies (15, 18, 23, 62). Mosaic 8b nanoparticles produce somewhat higher levels of memory antibodies with increased breadth and potency against heterologous viruses. However, SARS-CoV-2 mRNA vaccination elicits comparable serologic neutralizing responses after prime boost vaccination.

## Materials and Methods

### RBD expression and preparation of RBD-mi3 nanoparticles

RBD expression and preparation of SpyCatcher003-mi3 particles was done as previously described (19, 44).

### Phylogenetic tree

A sequence alignment of Sarbecovirus RBDs domains was built using Geneious software. A phylogenetic tree was calculated from the amino acid alignment using Geneious software. A table with the percentage of amino acid identity was also generated (Table S1). All the RBD sequences can be found in GenBank; RaTG13-CoV (GenBank QHR63300; S protein residues 319-541), SHC014-CoV (GenBank KC881005; residues 307-524), Rs4081-CoV (GenBank KY417143; S protein residues 310-515), pangolin17-CoV (GenBank QIA48632; residues 317-539), RmYN02- CoV (GSAID EPI_ISL_412977; residues 298-503), Rf1-CoV (GenBank DQ412042; residues 310- 515), WIV1-CoV (GenBank KF367457; residues 307-528), Yun11-CoV (GenBank JX993988; residues 310-515) and BM4831-CoV (GenBank NC014470; residues 310-530).

### Mouse immunization

Female mice (C57BL/6 and BALB/c) purchased from Jackson aged 6–12 weeks were housed at a temperature of 22 ºC and a humidity of 30–70% under a 12 h–12 h light–dark cycle with ad libitum access to food and water. Mice were primed with homotypic beta (SARS-CoV-2 B1.351 RBD) nanoparticles (n=5), mosaic 8b nanoparticles (n=10) or Spikevax Moderna mRNA vaccine (n=10). Mice were bled on days 14 and 28 after immunization and boosted after the second bleed. Lymph nodes were harvested on day 56 after immunization. Homotypic beta and mosaic 8b nanoparticles were injected intramuscularly. Intramuscular injections were performed using 1μg of mRNA vaccine (1μg of mRNA vaccine in 50μl of PBS + 50μl Addavax) or 5μg of nanoparticles in 50μl of PBS + 50μl of Addavax (Invivogen). The experiments were performed according to protocols approved by the Rockefeller University Institutional Animal Care and Use Committee (IACUC).

### Mouse sampling

Serum was collected with Goldenrod animal lancet by submandibular puncture at intermediary time points and cardiac puncture when the experiments were terminated in Serum gel Z1/1.1 (SARSTEDT), centrifuged at 1500rpm for 1 minute and stored at -20 ºC. Lymph nodes were dissected, mashed through a 70-μm cell strainer with 5ml of RPMI and the cells labeled for flow cytometry or frozen in fetal bovine serum (FBS) with 10% dimethyl sulfoxide in a gradual-freezing (∼1°C/min) container. Frozen cells were thawed in a 37°C water bath and immediately transferred to a prewarmed medium consisting of RPMI 1640, supplemented with 10% heat-inactivated FBS, 10 mM Hepes, 1× antibiotic-antimycotic, 1 mM sodium pyruvate, 2 mM l-glutamine, and 53 μM 2- mercaptoethanol (all from Gibco).

### ELISAs

We performed Enzyme-Linked Immunosorbent Assays (ELISAs) to evaluate plasma and monoclonal antibodies binding capacity to a panel of RBDs from SARS-CoV-2 (Wuhan-Hu-1), SARS-CoV-2 Omicron BA.1, SARS-CoV-2 Omicron XBB1.5, SARS-CoV-2 Delta (B.1.617), SARS-CoV-2 B1.351, Rs4081, BM4831, Yun11, SARS-CoV, Pang17, RatG13, Rf1, SHC014. 50 μl of a 1μg/ml solution of the relevant RBD in Phosphate-buffered Saline (PBS) was coated on high-binding 96-half-well plates (Corning 3690) overnight at 4C. Plates were washed 6 times with washing buffer (1× PBS with 0.05% Tween-20 (Sigma-Aldrich) and further incubated for 2 hours at room temperature with 170 μl of blocking buffer (1× PBS with 1% BSA and 0.05% Tween-20 (Sigma) and 0.1mM EDTA). Following the blocking step plasma samples or monoclonal antibodies were added in PBS and incubated for 1 hour at room temperature. Plasma samples were diluted (1:50) and serially diluted (1:3) 10 times. Monoclonal antibodies were used at 10 μg/ml starting concentration and serially diluted (1:4) 10 times. Plates were washed 6 times with washing buffer and incubated with secondary antibody (anti-human IgG for monoclonal and Anti-Mouse IgG for Plasma samples) conjugated to horseradish peroxidase (HRP) (Jackson Immuno Research 109-036-088 109-035-129 and Sigma A0295) in blocking buffer (1:5.000). Finally, plates were washed 6 times with washing buffer and further developed by addition of 50 μl of the HRP substrate, 3,3’,5,5’-Tetramethylbenzidine (TMB) (ThermoFisher) for 5 minutes. The reaction was stopped with 50 μl of 1 M H2SO4 solution and absorbance was measured at 450 nm with ELISA microplate reader (FluoStar Omega, BMG Labtech). Omega and Omega MARS software were used for the data analysis. A positive control monoclonal antibody C837 was used for anti-RBD ELISA and added to every plate for normalization for plasma samples. The average of its signal was used for normalization of all the other values on the same plate with Excel software. We determined the half-maximal binding titer or area under the curve for plasma (AUC) using four-parameter nonlinear regression in GraphPad Prism V9.1. 3BNC117, an HIV antibody, was used as a negative control for validation (55). For monoclonal antibodies, half-maximal concentration (EC50) was determined using four-parameter nonlinear regression in GraphPad Prism V9.1. The monoclonal antibodies with an EC50s above 1000 ng/mL were considered non-binders. All reported values represent the average of at least 3 independent experiments.

### SARS-CoV-2 pseudotyped reporter virus

The plasmids pSARS-CoV-SΔ19 and pSARS-CoV-2-SΔ19 (based on Wuhan-Hu-1 spike) expressing a C-terminally truncated SARS-CoV or SARS-CoV-2 spike protein, respectively, have been described (55, 63). Variant pseudoviruses resembling SARS-CoV-2 variants SARS-CoV-2 B.1.351 (11) and Omicron BA.1 have been described (64), as was a plasmid encoding for SARS-CoV-2 variant Omicron XBB1.5 (56). Plasmids were generated by introduction of substitutions using synthetic gene fragments (IDT) or overlap extension PCR mediated mutagenesis and Gibson assembly. Specifically, the variant-specific deletions and substitutions introduced were: Beta B.1.351: D80A, D215G, L242H, R246I, K417N, E484K, N501Y, D614G, A701V Omicron BA.1: A67V, Δ69-70, T95I, G142D, Δ143-145, Δ211, L212I, ins214EPE, G339D, S371L, S373P, S375F, K417N, N440K, G446S, S477N, T478K, E484A, Q493K, G496S, Q498R, N501Y, Y505H, T547K, D614G, H655Y, H679K, P681H, N764K, D796Y, N856K, Q954H, N969H, N969K, L981F Omicron XBB.1.5: T19I, L24S, del25-27, V83A, G142D, del144, H146Q, Q183E, V213E, G252V, G339H, R346T, L368I, S371F, S373P, S375F, T376A, D405N, R408S, K417N, N440K, V445P, G446S, N460K, S477N, T478K, E484A, F486P, F490S, Q498R, N501Y, Y505H, D614G, H655Y, N679K, P681H, N764K, D796Y, Q954H, N969K Deletions/substitutions corresponding to the variants listed above were incorporated into a spike protein that also includes the R683G substitution, which disrupts the furin cleavage site and increases particle infectivity. Neutralizing activity against mutant pseudoviruses were compared to a wildtype (WT) SARS-CoV-2 spike sequence (NC_045512), carrying R683G.

SARS-CoV-2 pseudotyped particles were generated as previously described (55, 63). Briefly, 293T (CRL-11268) cells were obtained from ATCC, and the cells were transfected with pNL4- 3ΔEnv-nanoluc and pSARS-CoV-2-SΔ19, particles were harvested 48 hours post-transfection, filtered, and stored at -80°C.

### Pseudotyped virus neutralization assay

Mouse sera, or monoclonal antibodies were five-fold serially diluted and incubated with SARS-CoV-2, SARS-CoV-2 B.1.351, SARS-CoV-2 BA.1, SARS-CoV-2 XBB1.5 or SARS-CoV pseudotyped virus for 1 hour at 37 °C. The mixture was subsequently incubated with HT1080/Ace2 cl14 cells for 48 hours after which cells were washed with PBS and lysed with Luciferase Cell Culture Lysis 5× reagent (Promega). Nanoluc Luciferase activity in lysates was measured using the Nano-Glo Luciferase Assay System (Promega) with the ClarioStar Microplate Multimode Reader (BMG). The relative luminescence units were normalized to those derived from cells infected with SARS-CoV-2, SARS-CoV-2 B.1.351, SARS-CoV-2 BA.1, SARS-CoV-2 XBB1.5 or SARS-CoV pseudotyped virus in the absence of serum or monoclonal antibodies. The half-maximal neutralization titers for serum (NT50) or half-maximal inhibitory concentrations for monoclonal antibodies (IC50) were determined using four-parameter nonlinear regression (least squares regression method without weighting; constraints: top=1, bottom=0) (GraphPad Prism9).

### Biotinylated protein for flow cytometry

SARS-CoV-2 RBD (Wuhan Hu-1) was produced as an Avi-tagged protein and biotinylated with the Biotin-Protein Ligase-BIRA kit following the manufacturer’s protocol (55). Ovalbumin (Sigma, A5503-1G) was biotinylated using the EZ-Link Sulfo-NHS-LC-Biotinylation kit according to the manufacturer’s instructions (Thermo Scientific). Biotinylated ovalbumin was conjugated to streptavidin-BV421 and used to remove unspecific binding for the single cell sorting experiments (Biolegend, 40-52-07)). SARS-CoV-2 RBD was conjugated to streptavidin-PE (BD Biosciences, 55-40-61) and streptavidin-APC (Biolegend 40-52-07) for single-cell sorting.

### Flow cytometry and single cell sorting

B cells were enriched by negative selection using a pan-B-cell isolation kit following the manufacturer’s instructions (Miltenyi Biotec, 130-101-638). The enriched B cells suspension were incubated in Flourescence-Activated Cell-sorting (FACS) buffer (1 x PBS, 2% FCS, 1 mM ethylenediaminetetraacetic acid (EDTA)) were washed and resuspended in a solution of mouse Fc-receptor blocking (BD pharmigen 1279893), fluorophore-conjugated bait (SARS-CoV-2 RBD-APC+, SARS-CoV-2 RBD-PE+) and Zombie-NIR Live/Dead cell marker for 15 min on ice. A master mix of anti-mouse antibodies (1:200) was used to label the cells for another 30 minutes incubation; anti-GL7-FITC (BD pharmigen, 55366), anti-CD38 AF700 (Invitrogen, 56038182) anti-CD95-PECy7 (BD Biosciences, 557653), anti-CD20-BV711 (BD horizon, 563832), anti-CD4- APC-eFluro 780 (Invitrogen, 47004282), anti-CD8-APC-eFluor 780 (Invitrogen, 47008182), anti- NKK1.1-APC-eFluor 780 (Invitrogen,47594182) and anti-F4/80 APC-eFluro 780 (Invitrogen, 47480182). Single B cell were sorted according to this phenotype CD4-, CD8-, NKK1.1-, GR1-, CD20+, CD95+, GL7+, Ova— RBD-PE+ and RBD-APC+ into individual wells of 96-well plates containing 4 μl of TE buffer + 2% β-Mercaptoethanol using FACS Aria III. For the acquisition FACSDiva software (Becton Dickinson) was used and FlowJo software for analysis. Sorted B cells were frozen on dry ice and stored at -80 °C.

### Antibody sequencing, cloning and expression

All monoclonal antibodies were identified and sequenced as described previously (55, 65). Following single cell sorting, RNA was cleaned up with RNAClean XP (Beckman Coulter, A63987) following the manufacturer’s instructions. Clean RNA from single cells was reverse transcribed with SuperScript III Reverse Transcriptase (Invitrogen, 18080-044). The cDNA was stored at −20 °C or used for nested PCR and Sanger sequencing of the variable IGH and IGK genes. Amplicons from the second PCR reaction (heavy and light chain) were sent for sequencing at Genewiz and further used as templates for sequence- and ligation-independent cloning into human antibody expression vectors. Recombinant monoclonal antibodies were produced and purified as described (55).

### Biolayer interferometry

We performed BLI experiments to measure antibody affinity with Octet Red (ForteBio) at 30 °C with shaking at 1,000 r.p.m. Epitope binding assays were performed with protein A biosensor (ForteBio 18-5010), following the manufacturer’s protocol: (1) Sensor check: sensors immersed 30 sec in buffer alone (buffer ForteBio 18-1105), (2) Capture 1st Ab: sensors immersed 10 min with Ab1 at 10 μg/mL, (3) Baseline: sensors immersed 30 sec in buffer alone, (4) Blocking: sensors immersed 5 min with IgG isotype control at 10 μg/mL. (5) Baseline: sensors immersed 30 sec in buffer alone, (6) Antigen association: sensors immersed 5 min with RBDs at 10 μg/mL. (7) Baseline: sensors immersed 30 sec in buffer alone. (8) Association Ab2: sensors immersed 5 min with Ab2 at 10 μg/mL. Curve fitting was performed using Fortebio Octet Data analysis software (ForteBio). Anti-RBD affinity measurements were corrected by subtracting the signal from IgGs in the absence of RBDs. The kinetic analysis using protein A biosensor was performed as follows: (1) baseline: 60sec immersion in buffer. (2) loading: 200sec immersion in a solution with IgGs 10 μg/ml. (3) baseline: 200sec immersion in buffer. (4) Association: 300sec immersion in solution with WT SARS-CoV-2 RBD at 20, 10 or 5 μg/ml (5) dissociation: 600sec immersion in buffer. Curve fitting was performed using a fast 1:1 binding model and Data analysis software (ForteBio). Mean KD values were determined by averaging all binding curves that matched the theoretical fit with an R2 value ≥ 0.8. All reported values represent the average of at least two independent experiments.

### Computational analyses of antibody sequences

Antibody sequences were trimmed based on quality and annotated using Igblastn v.1.14. with IMGT domain delineation system. Annotation was performed systematically using Change-O toolkit v.0.4.540 (66). Clonality of heavy and light chain was determined using DefineClones.py implemented by Change-O v0.4.5 (66). The script calculates the Hamming distance between each sequence in the data set and its nearest neighbor. Distances are subsequently normalized and to account for differences in junction sequence length, and clonality is determined based on a cut-off threshold of 0.15. Heavy and light chains derived from the same cell were subsequently paired, and clonotypes were assigned based on their V and J genes using in-house R and Perl scripts. All scripts and the data used to process antibody sequences are publicly available on GitHub (https://github.com/stratust/igpipeline/tree/igpipeline2_timepoint_v2).

The frequency distributions of mouse V genes in anti-SARS-CoV-2 antibodies from this study were compared to 131,284,220 IgH and IgL sequences generated by (67) and downloaded from cAb-Rep (68), a database of human shared BCR clonotypes available at https://cab-rep.c2b2.columbia.edu/. We selected the IgH and IgL sequences from the database that are partially coded by the same V genes and counted them according to the constant region. The frequencies shown in fig. S2 are relative to the source and isotype analyzed. We used the two-sided binomial test to check whether the number of sequences belonging to a specific IGHV or IGLV gene in the repertoire is different according to the frequency of the same IgV gene in the database. Adjusted p-values were calculated using the false discovery rate (FDR) correction.

Significant differences are displayed with numeric values on the graph.

Nucleotide somatic hypermutation and Complementarity-Determining Region (CDR3) length were determined using in-house R and Perl scripts. For somatic hypermutations, IGHV and IGLV nucleotide sequences were aligned against their closest germlines using Igblastn and the number of differences was considered to correspond to nucleotide mutations. The average number of mutations for V genes was calculated by dividing the sum of all nucleotide mutations across all participants by the number of sequences used for the analysis.

#### Data and materials availability

Figures and statistics were made with GraphPad Prism9 and arranged in Adobe Illustrator 2022. Comparisons between groups (homotypic beta, mosaic 8b, or mRNA) were calculated using an unpaired or paired t test in Prism 9.0 or multiple comparison (ANOVA). Differences were considered significant when P values were less than 0.05. Computer code to process the antibody sequences is available at GitHub (https://github.com/stratust/igpipeline/tree/igpipeline2_timepoint_v2). “All data are available in the main text or the supplementary materials.”

## Acknowledgments

We thank all members of the M.C.N. laboratory for helpful discussions, Jean-Philip Truman for technical assistance with cell-sorting experiments. This work was supported in part by National Institutes of Health (NIH) grant 5R37 AI037526, NIH Center for HIV/AIDS Vaccine Immunology and Immunogen Discovery (CHAVID) 1UM1AI144462-01 to M.C.N, and the Stavros Niarchos Foundation Institute for Global Infectious Disease Research. The work was also supported in part by Division of Intramural Research Program NIAD/NIH. M.C.N. is a Howard Hughes Medical Institute (HHMI) investigator. Furthermore we would like to thank Pamela Bjorkman and Jennifer Keeffe for sharing homotypic beta and Mosaic 8b nanoparticles with us.

## Figures and Tables

**Table 1. Sequences of anti-SARS-CoV-2 RBD IgG antibodies.** Sequences derived from each mouse are shown separately for homotypic beta, mosaic 8b, and mRNA. Clonally related sequences (same V and J genes for heavy and light chains and similar CDR3) share the same background color; white background indicates singlets.

**Table 2. Sequences of monoclonal IgG antibodies produced.** First column of the table is the name of the antibody produced followed by the nucleotide sequence of the paired heavy in blue and light chain in white.

