## Supplemental data for "Memory B cell Development in Response to mRNA SARS-CoV-2 and Nanoparticle Immunization in Mice"

### Supporting Information for
### Memory B cell Development in Response to mRNA SARS-CoV-2 and Nanoparticle Immunization in Mice

Authors: Marie Wiatr1, Zijun Wang1, Marie Canis2, Brianna Hernandez1, Anna Gazumyan1, Gabriela S. Silva Santos1, Paul D. Bieniasz2,3, Theodora Hatziioannou2, Frauke Muecksch2,4*, Michel C. Nussenzweig1,3*

1Laboratory of Molecular Immunology, The Rockefeller University; New York, NY 10065, USA.
2Laboratory of Retrovirology, The Rockefeller University; New York, NY 10065, USA.
3 Howard Hughes Medical Institute, The Rockefeller University, New York, NY, USA
4Department of Infectious Diseases, Virology, Medical Faculty Heidelberg, Heidelberg University; 69120 Heidelberg, Germany

* Corresponding authors:
Michel Nussenzweig
1230 York Avenue, New York, NY 10065
(212)327-8067

Frauke Muecksch
Im Neuenheimer Feld 344, 69120 Heidelberg, Germany
+49(0)6221-5635643

**This PDF file includes:**

Figures S1 to S4
Tables S1

#### Figures

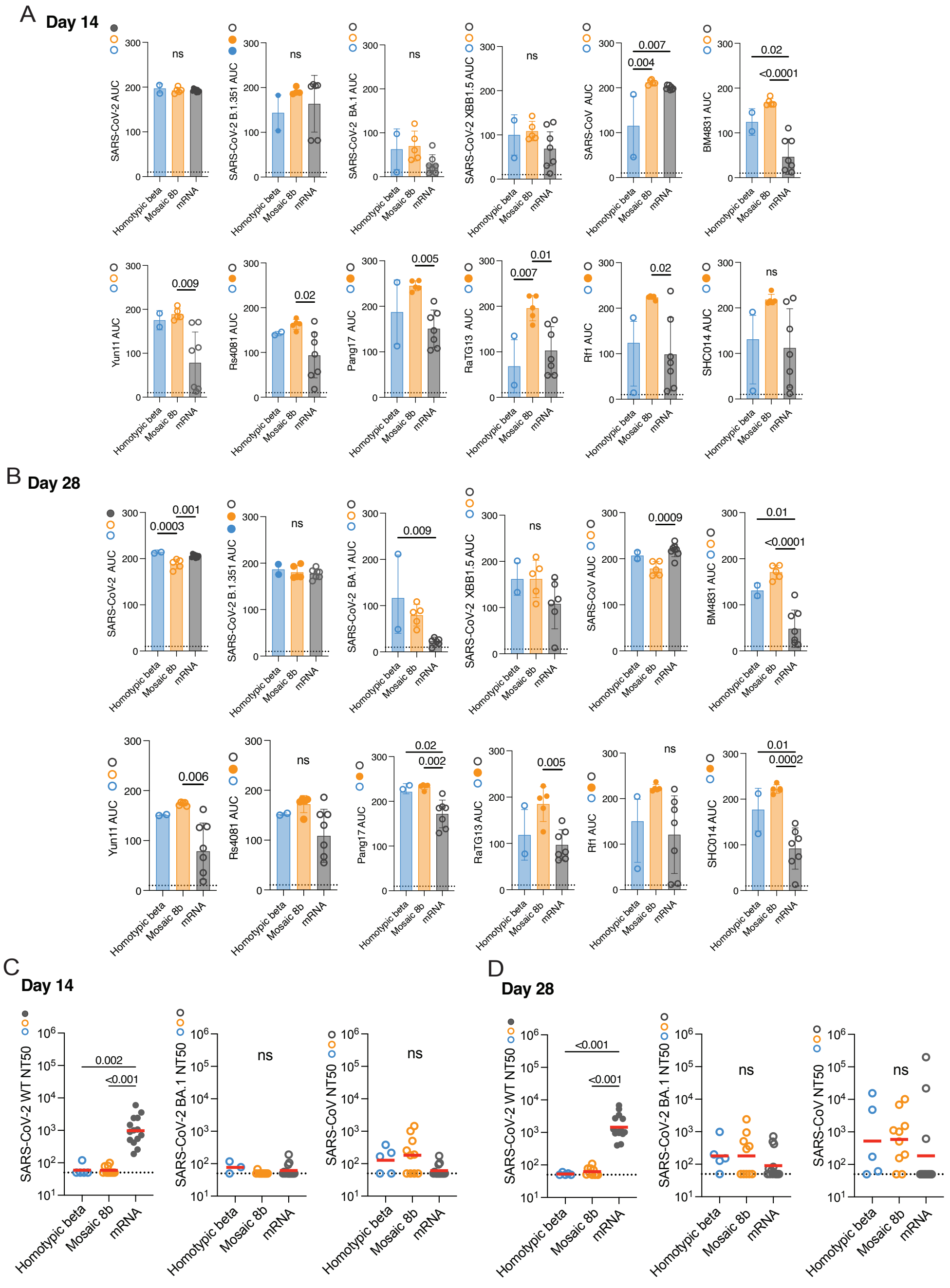


**Fig. S1. Plasma binding and neutralizing activity.** (A) Graphs show plasma IgG binding activity as normalized area under the curve (AUC) values measured by ELISA for; SARS-CoV-2, SARS-CoV-2 B.1.351, SARS-CoV-2 BA.1, SARS-CoV-2 XBB1.5, BM4831, Yun 11, Rs4081, Pang17, Ratg13, Rf1, and SHC014 RBDs on day 14 post prime. (B) Same as in (A) but measured on day 28. In (A) and (B) the dotted black line represents negative control. Each dot represents one animal. P values were calculated using an ANOVA test. (C and D) Plasma neutralizing titers for SARS-CoV-2, SARS-CoV-2 BA.1 and SARS-CoV. Plasma NT50s for homotypic beta (blue), mosaic 8b (orange) and mRNA (gray) measured on day 14 (C) and day 28 (D). Each dot represents one animal. The horizontal bar represents the geometric mean. All experiments were performed at least 2X. P values were calculated using Kruskal-Wallis test followed by Dunn’s multiple comparison test. Only P-values lower than 0.05 are shown.

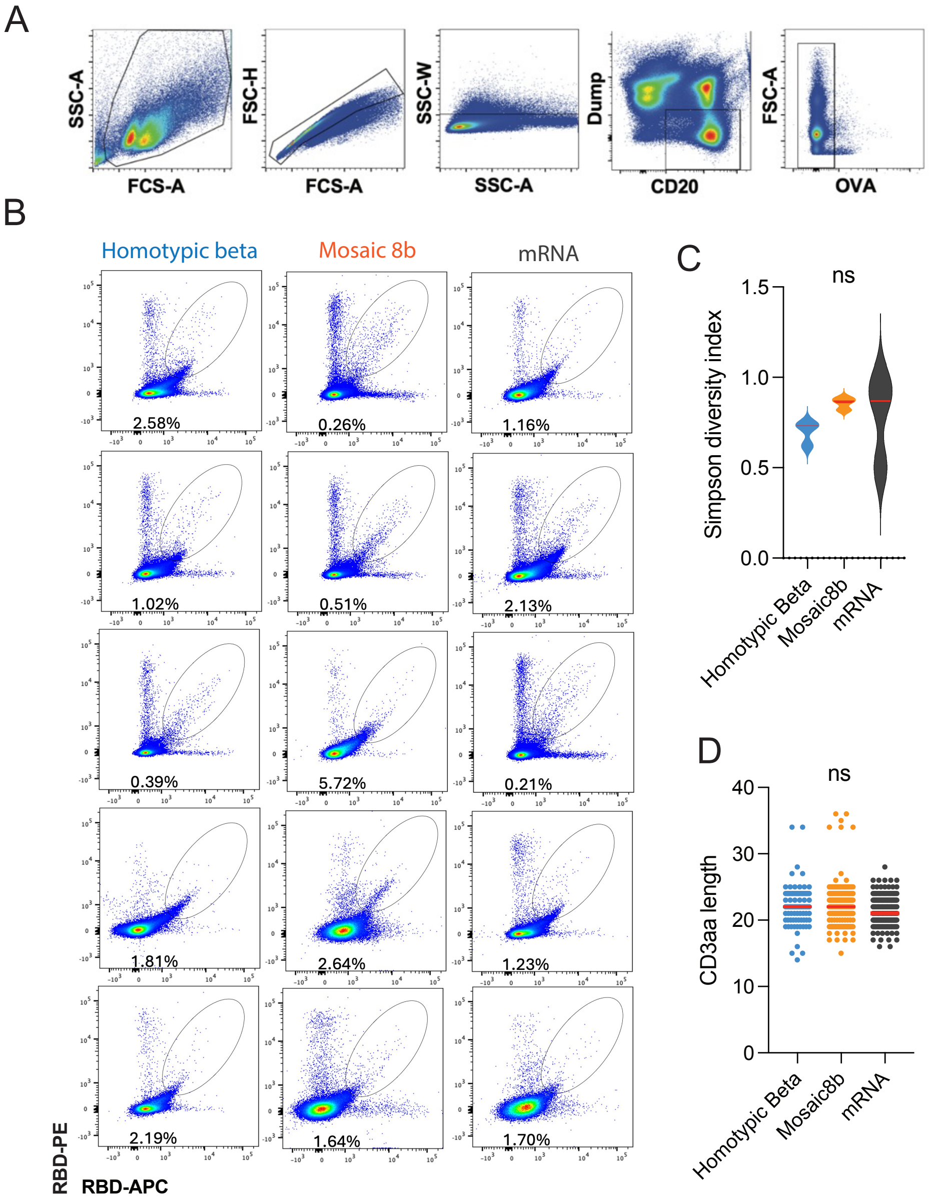


**Fig. S2. Flow cytometry.** (A) Gating strategy used for cell sorting. Doublets were excluded, CD20+ B cells selected and CD4+, CD8+, NKK1.1+, GR1+, OVA+ cells excluded. (B) Flow cytometry plots show the percentage of RBD+ B cells binding to phycoerythrin and allophycocyanin labeled RBDs in individual mouse lymph node on day 56 after immunization for homotypic beta-, mosaic 8b-, and mRNA-vaccinated mice. (C) Simpson diversity index for all antibodies in Fig. 2B and Table 1. (D) Combined CDR3 length for IgH and IgL for all antibodies in Fig. 2B and Table 1.

A

| ID | *Homotypic beta* SARS-CoV-2 WT | SARS-CoV-2 B1.351 | SARS-CoV-2 BA.1 | SARS-CoV-2 XBB1.5 | SARS-CoV | Epitope |
|---|---|---|---|---|---|---|
| 752 | 1.2 | 0.3 | 2.6 | 1000.0 | 1000.0 | II/III |
| 754 | 2.6 | 0.3 | 7.4 | 1000.0 | 1000.0 | II/III |
| 755 | 5.4 | 0.3 | 21.5 | 885.7 | 1000.0 | II/III |
| 757 | 6.9 | 0.3 | 16.8 | 952.6 | 1000.0 | II/III |
| 770 | 7.5 | 0.7 | 7.1 | 1000.0 | 1000.0 | II/III |
| 758 | 9.0 | 1.0 | 21.4 | 1000.0 | 1000.0 | II/III |
| 759 | 9.5 | 0.8 | 11.7 | 1000.0 | 1000.0 | II/III |
| 750 | 10.7 | 0.5 | 22.8 | 27.5 | 141.2 | II/III |
| 760 | 12.5 | 1.3 | 10.3 | 1000.0 | 1000.0 | II/III |
| 751 | 35.0 | 1.4 | 52.4 | 1000.0 | 1000.0 | II/III |
| 676 | 140.3 | 1000.0 | 733.8 | 1000.0 | 150.3 | UN |
| 625 | 262.9 | 1000.0 | 744.5 | 456.0 | 1000.0 | I |
| 769 | 307.9 | 0.6 | 629.4 | 63.5 | 313.5 | I/II/III/IV |
| 673 | 343.9 | 1000.0 | 1000.0 | 641.7 | 1000.0 | IV |
| 675 | 373.8 | 30.4 | 1000.0 | 1000.0 | 1000.0 | V |
| 687 | 422.2 | 1000.0 | 1000.0 | 1000.0 | 1000.0 | UN |
| 762 | 855.7 | 439.1 | 1000.0 | 1000.0 | 44.1 | I |
| 624 | 1000.0 | 632.4 | 1000.0 | 1000.0 | 1000.0 | UN |
| 674 | 1000.0 | 880.6 | 1000.0 | 1000.0 | 1000.0 | IV |
| 626 | 1000.0 | 1000.0 | 1000.0 | 1000.0 | 1000.0 | V |
| 627 | 1000.0 | 1000.0 | 1000.0 | 1000.0 | 1000.0 | I/II/III/IV |
| 628 | 1000.0 | 1000.0 | 1000.0 | 1000.0 | 1000.0 | UN |
| 629 | 1000.0 | 1000.0 | 1000.0 | 1000.0 | 1000.0 | I |
| 630 | 1000.0 | 1000.0 | 1000.0 | 1000.0 | 1000.0 | I/II/III/IV |
| 670 | 1000.0 | 1000.0 | 1000.0 | 1000.0 | 1000.0 | I/II |
| 671 | 1000.0 | 1000.0 | 1000.0 | 1000.0 | 1000.0 | I |
| 672 | 1000.0 | 1000.0 | 1000.0 | 1000.0 | 1000.0 | UN |
| 677 | 1000.0 | 1000.0 | 1000.0 | 1000.0 | 1000.0 | V |
| 681 | 1000.0 | 1000.0 | 1000.0 | 1000.0 | 1000.0 | UN |
| 685 | 1000.0 | 1000.0 | 1000.0 | 1000.0 | 1000.0 | UN |
| 768 | 1000.0 | 1000.0 | 1000.0 | 1000.0 | 1000.0 | I/III/IV |

| ID | *Mosaic 8b* SARS-CoV-2 WT | SARS-CoV-2 B1.351 | SARS-CoV-2 BA.1 | SARS-CoV-2 XBB1.5 | SARS-CoV | Epitope |
|---|---|---|---|---|---|---|
| 734 | 3.5 | 0.3 | 4.4 | n.d. | n.d. | I |
| 728 | 4.7 | 0.3 | 2.0 | n.d. | 750.0 | I/II |
| 740 | 17.2 | 0.3 | 144.3 | 0.3 | 16.8 | I/V |
| 716 | 54.8 | 0.3 | 2.9 | 138.2 | 1.1 | I/V |
| 711 | 64.3 | 0.6 | 2.5 | 240.8 | 1.1 | I |
| 745 | 74.1 | 178.3 | 57.8 | 61.3 | 1000.0 | UN |
| 721 | 119.1 | 0.3 | 1000.0 | 43.7 | 22.3 | IV/V |
| 713 | 135.2 | 0.3 | 6.3 | 8.7 | 4.8 | I |
| 735 | 189.1 | 0.3 | 603.7 | 2.4 | 62.4 | I/V |
| 718 | 208.1 | 30.8 | 1000.0 | 169.7 | 16.6 | IV/V |
| 702 | 213.0 | 0.3 | 656.6 | 2.1 | 13.9 | I/V |
| 739 | 233.6 | 0.3 | 783.0 | 13.6 | 122.6 | I |
| 729 | 340.9 | 50.3 | 212.5 | 1000.0 | 8.2 | I |
| 602 | 412.7 | 137.6 | 214.6 | 864.7 | 1000.0 | UN |
| 719 | 466.2 | 25.6 | 1000.0 | 1000.0 | 741.8 | IV/V |
| 710 | 651.6 | 2.3 | 18.9 | 1000.0 | 6.5 | I/II/III/IV |
| 703 | 687.7 | 0.3 | 746.0 | 61.7 | 733.6 | UN |
| 707 | 744.0 | 0.3 | 1000.0 | 839.0 | 1000.0 | UN |
| 404 | 826.3 | 1000.0 | 1000.0 | 1000.0 | 8.8 | UN |
| 725 | 848.7 | 2.3 | 1000.0 | 1000.0 | 164.0 | IV/V |
| 732 | 854.7 | 0.3 | 958.0 | 1000.0 | 1000.0 | I/II/III/IV |
| 619 | 905.0 | 415.2 | 1000.0 | 1000.0 | 6.8 | I/IV |
| 420 | 908.4 | 1000.0 | 1000.0 | 1000.0 | n.d. | UN |
| 705 | 1000.0 | 0.3 | 276.7 | 864.0 | 1000.0 | UN |
| 706 | 1000.0 | 0.3 | 923.1 | 1000.0 | 1000.0 | UN |
| 704 | 1000.0 | 0.3 | 1000.0 | 1000.0 | 1000.0 | UN |
| 731 | 1000.0 | 1.2 | 1000.0 | 1000.0 | 740.6 | I |
| 736 | 1000.0 | 2.5 | 1000.0 | n.d. | n.d. | IV |
| 724 | 1000.0 | 30.5 | 563.9 | 1000.0 | 140.5 | IV/V |
| 714 | 1000.0 | 37.0 | 68.3 | 1000.0 | 207.8 | IV/V |
| 726 | 1000.0 | 301.6 | 1000.0 | 1000.0 | 1000.0 | III |
| 742 | 1000.0 | 681.8 | 1000.0 | n.d. | 1000.0 | I/II/III |
| 401 | 1000.0 | 855.3 | 1000.0 | 1000.0 | 1000.0 | I/IV |
| 744 | 1000.0 | 1000.0 | 1000.0 | 810.1 | 1000.0 | UN |
| 730 | 1000.0 | 1000.0 | 1000.0 | 1000.0 | 20.3 | I |
| 400 | 1000.0 | 1000.0 | 1000.0 | 1000.0 | 1000.0 | I/IV |
| 403 | 1000.0 | 1000.0 | 1000.0 | 1000.0 | 1000.0 | IV |
| 407 | 1000.0 | 1000.0 | 1000.0 | 1000.0 | 1000.0 | UN |
| 601 | 1000.0 | 1000.0 | 1000.0 | 1000.0 | 1000.0 | I |
| 606 | 1000.0 | 1000.0 | 1000.0 | 1000.0 | 1000.0 | I |
| 620 | 1000.0 | 1000.0 | 1000.0 | 1000.0 | 1000.0 | I/IV |
| 727 | 1000.0 | 1000.0 | 1000.0 | 1000.0 | 1000.0 | I |
| 607 | 1000.0 | 1000.0 | 1000.0 | 1000.0 | n.d. | IV |
| 746 | 1000.0 | 1000.0 | 1000.0 | 1000.0 | 1000.0 | I/IV |
| 733 | 1000.0 | 30.9 | 1000.0 | 947.1 | 1000.0 | V |
| 723 | 1000.0 | n.d. | 1000.0 | 612.1 | 817.8 | I/IV |
| 722 | 1000.0 | 1000.0 | 1000.0 | 1000.0 | 1000.0 | V |

| ID | *mRNA* SARS-CoV-2 WT | SARS-CoV-2 B1.351 | SARS-CoV-2 BA.1 | SARS-CoV-2 XBB1.5 | SARS-CoV | Epitope |
|---|---|---|---|---|---|---|
| 900 | 0.5 | 0.3 | 780.5 | 1000.0 | 1000.0 | III |
| 512 | 0.6 | 0.3 | 1000.0 | 1000.0 | 1000.0 | III |
| 323 | 0.8 | 0.3 | 1000.0 | 1000.0 | 1000.0 | UN |
| 332 | 1.1 | 331.2 | 1000.0 | 1000.0 | 1000.0 | III |
| 327 | 1.1 | 0.3 | 1000.0 | 1000.0 | 1000.0 | UN |
| 330 | 1.2 | 0.3 | 1000.0 | 1000.0 | 1000.0 | III |
| 329 | 1.5 | 0.3 | 1000.0 | 1000.0 | 1000.0 | UN |
| 331 | 1.5 | 0.3 | 1000.0 | 1000.0 | 1000.0 | III |
| 335 | 2.1 | 0.3 | 1000.0 | 1000.0 | 1000.0 | III |
| 328 | 2.3 | 0.3 | 274.0 | 1000.0 | 1000.0 | UN |
| 909 | 8.6 | 1.7 | n.d. | 48.3 | 44.9 | I |
| 315 | 9.1 | 1000.0 | 1000.0 | 1000.0 | 1000.0 | UN |
| 326 | 9.4 | 0.3 | 1000.0 | 1000.0 | 1000.0 | UN |
| 316 | 16.3 | 1000.0 | 1000.0 | 1000.0 | 1000.0 | UN |
| 908 | 21.5 | 1000.0 | 1000.0 | 1000.0 | 1000.0 | UN |
| 913 | 36.6 | 2.5 | 42.6 | 93.7 | 98.3 | IV/V |
| 513 | 38.4 | 1000.0 | 873.3 | 1000.0 | 797.1 | II |
| 912 | 150.9 | 51.7 | 63.1 | 74.7 | 1000.0 | V |
| 917 | 173.4 | 1000.0 | 1000.0 | 1000.0 | 1000.0 | I/II |
| 341 | 178.8 | 64.7 | 204.9 | 1000.0 | 1000.0 | V |
| 349 | 203.6 | 43.7 | 157.0 | 219.2 | 1000.0 | V |
| 344 | 248.0 | 72.2 | 192.7 | 113.0 | 1000.0 | V |
| 804 | 349.4 | 700.9 | 1000.0 | 1000.0 | 1000.0 | IV |
| 340 | 549.5 | 91.1 | 142.0 | 314.9 | 1000.0 | V |
| 802 | 560.4 | 1000.0 | 1000.0 | 1000.0 | 1000.0 | IV |
| 343 | 594.6 | 244.2 | 91.1 | 426.6 | 1000.0 | V |
| 342 | 676.5 | 97.6 | 154.1 | 408.7 | 1000.0 | V |
| 348 | 727.9 | 88.5 | 104.6 | 89.4 | 1000.0 | V |
| 916 | 742.2 | 1000.0 | 1000.0 | 1000.0 | 1000.0 | I |
| 920 | 758.1 | 183.8 | 1000.0 | 1000.0 | 1000.0 | IV |
| 347 | 787.4 | 95.3 | 148.6 | 72.0 | 1000.0 | V |
| 806 | 864.3 | 1000.0 | 1000.0 | 1000.0 | 1000.0 | IV |
| 346 | 884.7 | 104.2 | 1000.0 | 1000.0 | 100.0 | IV |
| 907 | 911.2 | 1000.0 | 1000.0 | 1000.0 | 1000.0 | IV |
| 915 | 927.9 | 1000.0 | 1000.0 | 1000.0 | 1000.0 | IV |
| 805 | 1000.0 | 41.5 | 1000.0 | 1000.0 | 1000.0 | IV |
| 320 | 1000.0 | 186.6 | 1000.0 | 1000.0 | 1000.0 | V |
| 801 | 1000.0 | 661.2 | 1000.0 | 1000.0 | 1000.0 | IV |
| 345 | 1000.0 | 1000.0 | 1000.0 | 1000.0 | 200.1 | IV |
| 300 | 1000.0 | 1000.0 | 1000.0 | 1000.0 | 1000.0 | IV |
| 301 | 1000.0 | 1000.0 | 1000.0 | 1000.0 | 1000.0 | IV |
| 302 | 1000.0 | 1000.0 | 1000.0 | 1000.0 | 1000.0 | IV |
| 303 | 1000.0 | 1000.0 | 1000.0 | 1000.0 | 1000.0 | IV |
| 304 | 1000.0 | 1000.0 | 1000.0 | 1000.0 | 1000.0 | IV |
| 306 | 1000.0 | 1000.0 | 1000.0 | 1000.0 | 1000.0 | UN |
| 307 | 1000.0 | 1000.0 | 1000.0 | 1000.0 | 1000.0 | UN |
| 308 | 1000.0 | 1000.0 | 1000.0 | 1000.0 | 1000.0 | UN |
| 309 | 1000.0 | 1000.0 | 1000.0 | 1000.0 | 1000.0 | UN |
| 313 | 1000.0 | 1000.0 | 1000.0 | 1000.0 | 1000.0 | IV |
| 317 | 1000.0 | 1000.0 | 1000.0 | 1000.0 | 1000.0 | UN |
| 318 | 1000.0 | 1000.0 | 1000.0 | 1000.0 | 1000.0 | UN |
| 319 | 1000.0 | 1000.0 | 1000.0 | 1000.0 | 1000.0 | V |
| 321 | 1000.0 | 1000.0 | 1000.0 | 1000.0 | 1000.0 | IV |
| 352 | 1000.0 | 1000.0 | 1000.0 | 1000.0 | 1000.0 | IV |
| 355 | 1000.0 | 1000.0 | 1000.0 | 1000.0 | 1000.0 | UN |
| 356 | 1000.0 | 1000.0 | 1000.0 | 1000.0 | 1000.0 | V |
| 357 | 1000.0 | 1000.0 | 1000.0 | 1000.0 | 1000.0 | UN |
| 358 | 1000.0 | 1000.0 | 1000.0 | 1000.0 | 1000.0 | I |
| 819 | 1000.0 | 1000.0 | 1000.0 | 1000.0 | 1000.0 | I/II/III/IV |
| 820 | 1000.0 | 1000.0 | 1000.0 | 1000.0 | 1000.0 | I/II/III/IV |
| 902 | 1000.0 | 971.7 | 1000.0 | 1000.0 | 1000.0 | UN |
| 911 | 1000.0 | 1000.0 | 1000.0 | 1000.0 | 1000.0 | I |
| 914 | 1000.0 | 1000.0 | 1000.0 | 1000.0 | 1000.0 | II |
| 904 | n.d. | 10.6 | n.d. | n.d. | n.d. | V |
| 514 | n.d. | 689.9 | 1000.0 | 1000.0 | 1000.0 | UN |
| 800 | n.d. | 822.5 | 1000.0 | 787.3 | 722.9 | IV |

B

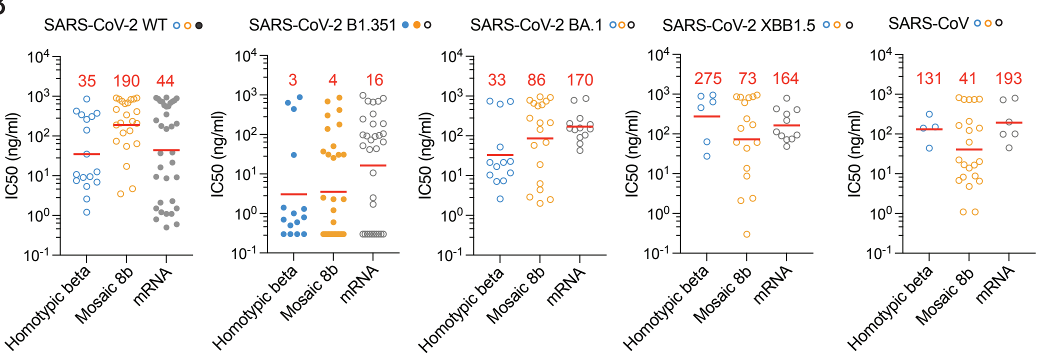


**Fig. S3. Neutralization activity of monoclonal antibodies.** (A) Tables showing the IC50 (ng/ml) for each monoclonal antibody against the 5 tested pseudoviruses, SARS-CoV-2, SARS-CoV-2 B.1.351, SARS-CoV-2 BA.1, SARS-CoV-2 XBB1.5, and SARS-CoV. Color coding is white=most potent neutralizers [IC50<10ng/ml], red=no neutralization [IC50>=1000ng/ml]. (B) SARS-CoV-2, SARS-CoV-2 B.1.351, SARS-CoV-2 BA.1, SARS-CoV-2 XBB1.5 and SARS-CoV pseudovirus neutralization activity for antibodies with a detectable IC50 values in ng/ml for the indicated pseudoviruses. The red line shows the geometric mean indicated in red. Statistical significance was determined using the Kruskal-Wallis test with subsequent Dunn's multiple-comparisons test.

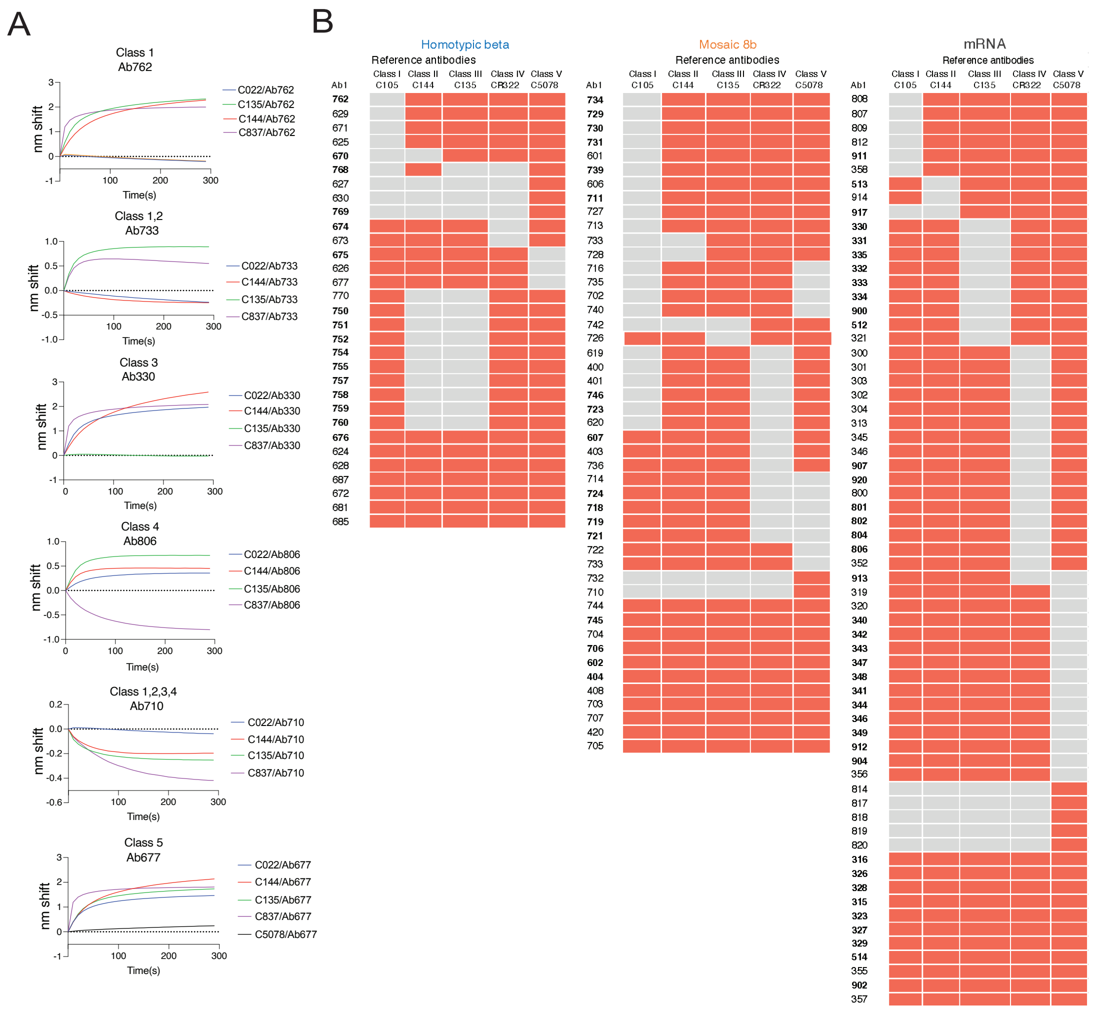


**Fig. S4. Biolayer interferometry determination of epitope binding.** (A) BLI curves showing interference of pre-formed antibody-RBD complexes with reference antibodies belonging to class 1 (C022, blue), class 2 (C144, yellow), class 3 (C135, orange), class 4 (C837, green) and class 5 (C5078, purple). (B) Tables display the BLI-mapped class of monoclonal antibodies for each immunization (homotypic beta, mosaic 8b and mRNA). Reference class-indicating antibodies are listed on the top. Grey indicates the class in which tested monoclonal antibodies belong, red the exclusion of the class. Bold antibodies are neutralizers.

#### Tables

| | SARS-CoV-2 | SARS-CoV-2 BA.1 | SARS-CoV-2 B1.351 | SARS-CoV-2 XBB1.5 | SARS-CoV-2 B1.617 | RaTG13 | SHC014 | Rs4081 | Rf1 | WIV1 | Yun11 | Pang17 | SARS-CoV | BM4831 | RmYN02 |
|---|---|---|---|---|---|---|---|---|---|---|---|---|---|---|---|
| SARS-CoV-2 | | 93.3 | 98.7 | 90.1 | 99.1 | 90.1 | 76.2 | 63.7 | 64.6 | 75.3 | 63.7 | 86.5 | 73.1 | 70.1 | 63.2 |
| SARS-CoV-2 BA.1 | 93.3 | | 94.2 | 94.2 | 93.3 | 87.4 | 75.3 | 62.8 | 63.7 | 72.2 | 61.9 | 83.0 | 70.9 | 68.3 | 61.9 |
| SARS-CoV-2 B1.351 | 98.7 | 94.2 | | 91.0 | 97.8 | 89.7 | 76.2 | 63.7 | 64.6 | 74.9 | 63.7 | 86.5 | 73.1 | 70.1 | 63.2 |
| SARS-CoV-2 XBB1.5 | 90.1 | 94.2 | 91.0 | | 90.1 | 86.1 | 74.9 | 61.0 | 62.3 | 71.3 | 60.1 | 82.1 | 70.0 | 67.4 | 60.1 |
| SARS-CoV-2 B1.617 | 99.1 | 93.3 | 97.8 | 90.1 | | 90.1 | 75.8 | 63.7 | 64.6 | 75.8 | 63.7 | 85.7 | 73.5 | 70.1 | 63.2 |
| RaTG13 | 90.1 | 87.4 | 89.7 | 86.1 | 90.1 | | 75.8 | 63.7 | 63.7 | 76.2 | 63.2 | 87.9 | 74.9 | 68.8 | 62.8 |
| SHC014 | 76.2 | 75.3 | 76.2 | 74.9 | 75.8 | 75.8 | | 63.1 | 64.4 | 84.2 | 64.4 | 75.8 | 82.0 | 74.9 | 64.0 |
| Rs4081 | 63.7 | 62.8 | 63.7 | 61.0 | 63.7 | 63.7 | 63.1 | | 89.7 | 64.9 | 92.6 | 65.0 | 63.5 | 62.1 | 88.7 |
| Rf1 | 64.6 | 63.7 | 64.6 | 62.3 | 64.6 | 63.7 | 64.4 | 89.7 | | 65.3 | 89.2 | 65.0 | 64.0 | 64.4 | 90.2 |
| WIV1 | 75.3 | 72.2 | 74.9 | 71.3 | 75.8 | 76.2 | 84.2 | 64.9 | 65.3 | | 66.2 | 74.9 | 94.6 | 73.5 | 64.9 |
| Yun11 | 63.7 | 61.9 | 63.7 | 60.1 | 63.7 | 63.2 | 64.4 | 92.6 | 89.2 | 66.2 | | 64.1 | 64.9 | 62.1 | 88.7 |
| Pang17 | 86.5 | 83.0 | 86.5 | 82.1 | 85.7 | 87.9 | 75.8 | 65.0 | 65.0 | 74.9 | 64.1 | | 74.9 | 68.3 | 63.2 |
| SARS-CoV | 73.1 | 70.9 | 73.1 | 70.0 | 73.5 | 74.9 | 82.0 | 63.5 | 64.0 | 94.6 | 64.9 | 74.9 | | 72.2 | 63.5 |
| BM4831 | 70.1 | 68.3 | 70.1 | 67.4 | 70.1 | 68.8 | 74.9 | 62.1 | 64.4 | 73.5 | 62.1 | 68.3 | 72.2 | | 62.1 |
| RmYN02 | 63.2 | 61.9 | 63.2 | 60.1 | 63.2 | 62.8 | 64.0 | 88.7 | 90.2 | 64.9 | 88.7 | 63.2 | 63.5 | 62.1 | |

**Table S1.** Percentage of amino acids similarity from the different Sarbecovirus RBD protein (receptor binding domain); RaTG13, SARS-CoV-2 B.1.351, Rs4081, SHC014, W1V1, BM4831, Rf1, SARS-CoV-2 B.1.617, SARS-CoV-2, SARS-CoV-2 BA.1, Yun11, SARS-CoV-2 XBB1.5, Pang17, RmYN02 and SARS-CoV. Color coding is white=low similarity between two RBDs, red= high similarity between two RBDs.
